# Ketone body metabolism declines with age in mice in a sex-dependent manner

**DOI:** 10.1101/2022.10.05.511032

**Authors:** Brenda Eap, Mitsunori Nomura, Oishika Panda, Thelma Y Garcia, Christina D King, Jacob P Rose, Teresa C Leone, Daniel P Kelly, Birgit Schilling, John C Newman

## Abstract

Understanding how our cells maintain energy homeostasis has long been a focus of aging biology. A decline in energy metabolism is central to many age-related diseases such as Alzheimer’s disease, heart failure, frailty, and delirium. Intervening on pathways involved in energy homeostasis can extend healthy lifespan. When the primary energy substrate glucose is scarce, mice and humans can partially switch cellular energetic needs to fat-derived ketone bodies (i.e., beta-hydroxybutyrate (BHB), acetoacetate, acetone). Aging is associated with glucose intolerance and insulin insensitivity, yet, surprisingly, what role ketone body metabolism might play in compensating for impaired glucose utilization in age-related diseases is understudied. Here, we investigate how endogenous ketone body production and utilization pathways are modulated by age across the lifespan of male and female C57BL/6N mice (3 mo old, 12 mo old, 22 mo old). We show how different ages have different metabolic and gene expression responses to 1-week ketogenic diet (KD). We hypothesized that there would be a compensatory ketogenic response with age but instead saw declines in plasma BHB concentrations under fasting and non-fasting conditions with strong sexual dimorphism. Under KD, both sexes increased BHB concentrations at all ages, but only males showed strong gene expression induction. We also observed tissue-specific changes with age in baseline ketone metabolism, and surprising induction of extrahepatic ketogenic genes under KD. We found significant residual blood concentrations of BHB in KD even after a knockout of liver BHB production. Overall, these findings show that older mice have impaired non-fasting ketogenesis but are capable of increasing their ketogenic capacity under stimulation (i.e., KD) to meet energetic demands in aging. Therapies to augment non-fasting ketogenesis or provide exogenous ketones may be useful to improve energy homeostasis in diseases and conditions of aging.

## Introduction

Aging is associated with glucose intolerance and insulin insensitivity, resulting in a deregulation of the nutrient-responsive pathway, a common hallmark of aging^1–3^. While lifestyle interventions like fasting, caloric restriction and dietary restriction can extend healthy mammalian lifespan, impactful translation to improve human health requires a deeper mechanistic understanding of these interventions. For examples, IGF-1 signaling and the mammalian target of rapamycin (mTOR) pathways are promising translational targets with drugs now in clinical trials (e.g., NCT02432287, NCT03733132, NCT01649960, and NCT05414292)^4–7^. One of the commonly observed metabolic change associated with these interventions across species is the elevation of blood ketone bodies. Similar to the discoveries of rapamycin and metformin, endogenous metabolites like ketone bodies have also been shown to prevent the deregulation of the nutrient-responsive pathway^8–13^.

Ketone bodies are synthesized in the liver from non-esterified fatty acids (NEFAs) mobilized from adipose tissue. Ketone bodies, of which beta-hydroxybutyrate (BHB) is most abundant, are commonly known as glucose-sparing energy molecules for extra-hepatic tissues such as brain, heart, kidney, and skeletal muscle^14^. In healthy adults, ketosis under non-fasting conditions maintains plasma BHB concentrations to a range of 50 to 250 μM^15^. However, a higher non-fasting ketosis can be achieved through fasting, exercise, and dietary interventions like a ketogenic diet, increasing concentrations above 500 μM and into the low millimolar range^16–22^. In addition to its role as an alternative oxidative fuel, BHB and acetoacetate are increasingly understood to have a variety of signaling activities also relevant to aging. BHB specifically has been shown to play important roles in health and disease. In brain aging, two studies showed that feeding a non-obese KD could extend the healthy lifespan of wild-type mice including improving memory function^20,21^. In humans, ketone body uptake differs in normal brain aging, mild cognitive impairment, and Alzheimer’s Disease^23^. KD, intermittent fasting, and ketogenic pharmacotherapies like SGLT2i have shown to protect against diabetic kidney disease^24^. In the failing heart, it’s been shown that the heart prefers ketone bodies as more energy efficient substrates, which helps explain why patients with heart failure have elevated concentrations of ketone bodies^25–27^. BHB has also shown to regulate inflammation by inhibiting the NLRP3 inflammasome, reducing proinflammatory IL-1B, IL-18, and caspase-1 activation, in heart, kidney and the central nervous system^28,29^.

This growing literature suggests that ketogenic interventions have promise for various diseases and conditions of aging, but there are significant gaps. Much of the work in aging was carried out in male mice only, despite clear metabolic differences between males and females in glucose and fat metabolism^30,31^. Work to date has focused on interventions, while how the baseline endogenous ketogenic system changes with age is not understood. And little is known about how the response of the endogenous system to interventions like KD changes with age. Our most recent work showed an improvement in memory of aging male mice, leading us to further elucidate the effects of KD across ages and sexes^20,21^. From the same work, we showed that ketogenesis is tightly regulated by the peroxisome proliferator activated receptor alpha (PPARα). Therefore, we sought to understand the regulation of ketone body metabolism across ages, sexes, and diets to investigate the hypothesis that ketogenesis is compensatory to aging and is tissue-specific to hypometabolism of glucose we commonly see with age. In this study, we first test the different responses of male and female C57BL/6 mice to fasting across different ages. We use a typical high-carbohydrate diet to study non-fasting ketogenesis across “normal” aging metabolism. We then use KD to stimulate the endogenous ketogenic system and test how the response differs across ages and sexes. We find that non-fasting hepatic ketogenesis declines across ages more so in males in both fasted and non-fasted states, but could be augmented by KD in a sex-dependent manner. We also show that aging induces organ-specific ketogenesis to compensate for age-related declines in energetic demands.

## Results

### Fasting-induced ketosis declines in aging and is sex-dependent

We first tested if the kinetics of fasting ketosis, particularly the production of BHB, changes with age, a topic surprisingly understudied given the links between aging biology and fasting^32^. Male and female C57BL/6N mice of three ages (young adult, 3 months old; middle-aged adult, 12 months old; and older adult, 22 months old) were fasted for 16 hours (6am-10pm) and monitored every 2 hours with blood drawn for glucose and BHB measurements. Both male and female blood glucose levels decreased to approximately 100 mg/dL by the end of their fasting period **(Fig. 1a, e)**. Blood BHB concentrations increased as expected, but with striking age- and sex-dependent variability (**Fig. 1b, f**). The overall area under the curve (AUC) of BHB concentration declined sharply with age in both sexes **(Fig. 1c, g)**. Interestingly, the trajectory of plasma BHB concentrations was different in females, with an early intermediate peak and faster late rise, compared to males. In sum, the overall BHB AUC during a 16-hour fast was 1.3-2 times higher in females compared to males, as was the final BHB concentration at the end of fasting **(Fig. 1d, h)**, despite the similar overall declines with age between sexes.

**Fig. 1.**
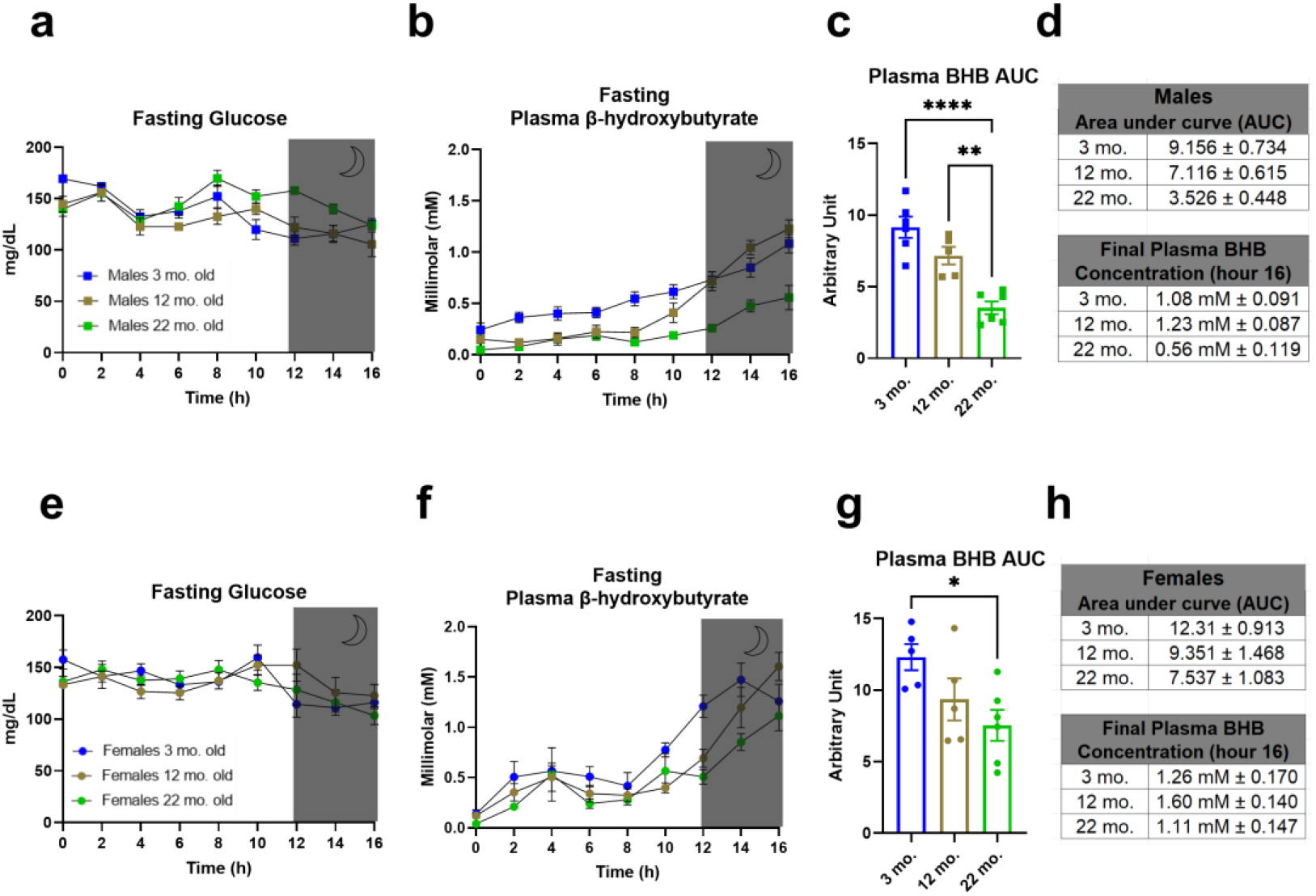
Fasting-induced ketosis declines in aging and is sex-dependent. **a**, Trajectory of 16-hour fasting glucose levels in male mice (3, 12, 22 months), 6am-10pm **b**, Trajectory of 16-hour fasting plasma BHB levels in males, 6am-10pm **c**, AUC of plasma BHB in males **d**, values of fasting AUC and final fasting concentration in males **e**, Trajectory of 16-hour fasting glucose levels in female mice (3, 12, 22 months) **f**, Trajectory of 16-hour fasting plasma BHB levels in females **g**, AUC of plasma BHB in females **h**, values of fasting AUC and final fasting concentration in females **a-c, e-g**, n=5-6 per age group, all data are presented as mean ± SEM; Ordinary one-way ANOVA, post-hoc Tukey’s for multiple comparisons test between age groups, *p<0.0332, **p<0.0021, ***p<0.0002, ****p<0.001.

### Age-related changes in metabolic state after fasting is sex-dependent

The fasting data suggest that both age and sex affect energy substrate plasticity. To better characterize differences in energy substrate utilization during fasting, we observed male and female C57BL/6N mice at the same three ages (3 mo, 12 mo, 22 mo) in metabolic chambers for 4.5 days with a 24-hour fasting period in between **(Fig. 2a, e)**. The respiratory exchange ratio (RER) is the ratio between CO_2_ emission and O_2_ consumption, with values typically ranging between 0.7 (primarily fat oxidation) and 1.0 (primarily carbohydrate oxidation). The RER of male mice both before and after fasting was around 0.94 at all ages **(Fig. 2b, d)**. In contrast, female mice at 12 and 22 months (but not 3 months) had a RER of 0.80 before fasting, increasing to 0.94 only after fasting with the smallest change in the oldest females (**Fig. 2f, h**). Therefore, older female mice appeared to be more dependent on fat oxidation at baseline compared to young female mice or male mice at all ages **(Extended Fig. 1a-c)**. During the fasting period, RER of all mice dropped to 0.79, a large change for males but minimal change from baseline for females (**Fig. 2c, g**). In contrast to the striking decline in fasting BHB with age, the differences with age in overall fuel utilization before, during, and after fasting via RER were relatively small. Fuel utilization did show important sex differences with fat metabolism predominating in non-fasting middle-aged and old female mice. This difference may begin to explain the consistently higher BHB concentrations in fasting observed for female mice.

**Fig. 2.**
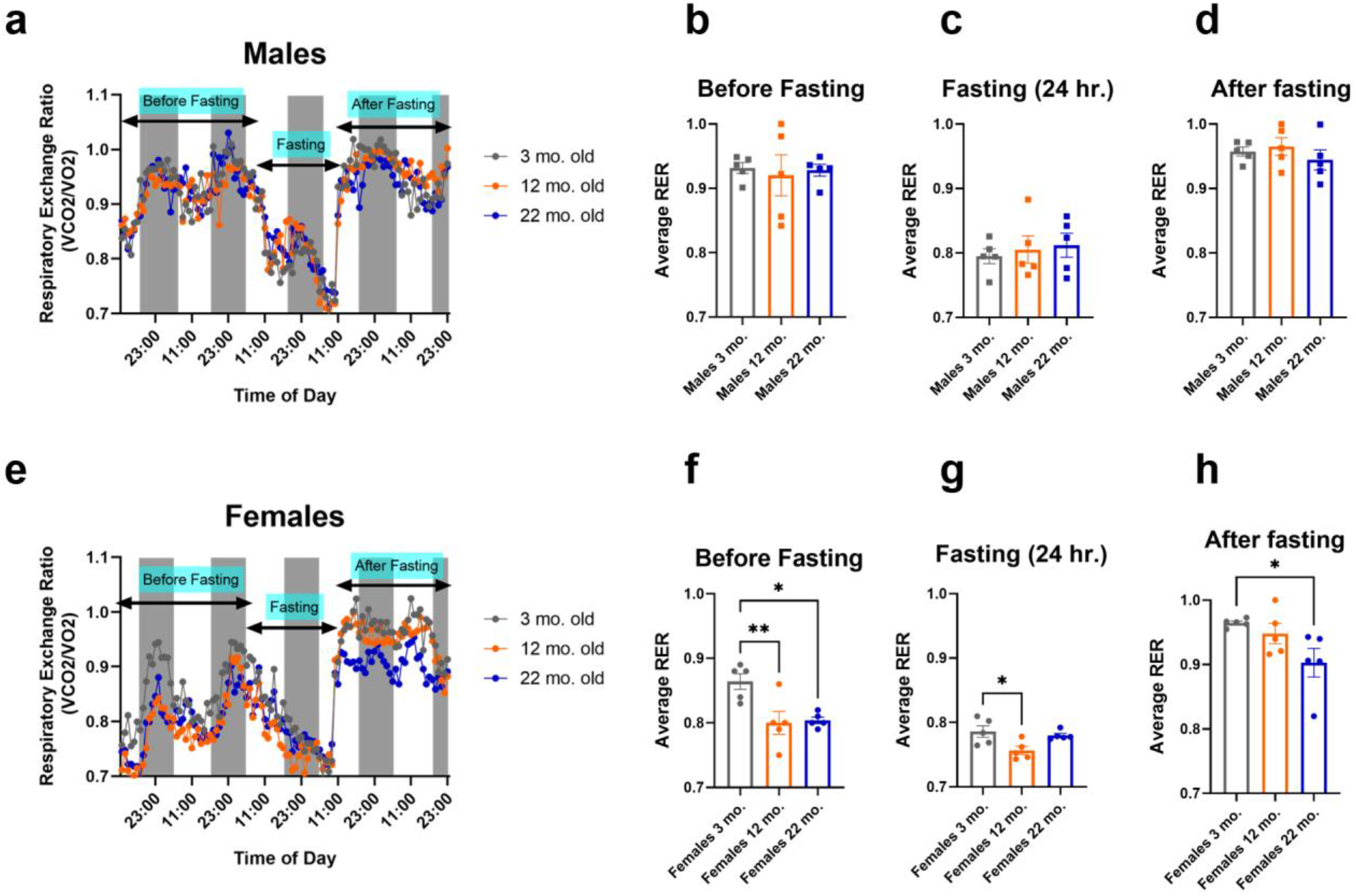
Age-related changes in metabolic state after fasting is sex-dependent. **a**, Respiratory Exchange Ratio (RER) of male mice (3, 12, 22 months) over the course of 4.5 days **b**, Average RER for before fasting in males **c**, Average RER for 24 hour fasting in males **d**, Average RER for after fasting in males **e**, Respiratory Exchange Ratio (RER) of female mice over the course of 4.5 days **f**, Average RER for before fasting in female mice (3, 12, 22 months) **g**, Average RER for 24 hour fasting in females **h**, Average RER for after fasting in females. **a**,**e**, n=5 per age group **b-d, f-h**, n=5 per age group; all data are presented as mean ± SEM; Ordinary one-way ANOVA, post-hoc Tukey’s for multiple comparisons test between age groups, *p<0.0332, **p<0.0021, ***p<0.0002. ****p<0.001.

### Ketogenic diet helps augment non-fasting ketosis in aging

To further understand how non-fasting ketosis changes in aging between sexes, the same three age groups of male and female mice were acclimated to a control diet (77% carbohydrate, 10% protein, 13% fat) for 14 days and then fed either the same control diet or a high-fat ketogenic diet (90% fat and 10% protein) for an additional week **(Fig. 3a, f)**. We collected tissue and plasma on day 7 just before the start of the light cycle (6am). A ketogenic diet is a means of stimulating ketone body production without fasting or energy restriction, and like fasting is a potent inducer of genes related to ketone body production and utilization^20^. We reasoned that the control-fed group would show how the basal ketogenic system changes with age, while the ketogenic diet would show the latent potential of the ketogenic system upon stimulation. Changes in body weights in both sexes on the control diet were less than a gram **(Extended Fig. 2a, b)**. Circulating plasma BHB concentrations were within the typical range (0.05-0.3 mM) of a predominantly carbohydrate diet and were similar between males and females, but were generally lower in older mice though this reached statistical significance only in females **(Fig. 3b)**. Two major regulators of ketogenesis are insulin and substrate availability in the form of non-esterified fatty acids (NEFA). Insulin levels increased with age as expected **(Fig. 3c)**, accompanied by a decline in NEFA **(Fig. 3d)** albeit again only significant in females. Glucose concentrations also declined with age in these non-obese mice **(Fig. 3e)**. Together the rise in insulin and decline in NEFA help explain the decline in basal, non-fasting BHB concentrations. Overall, these data show that non-fasting ketosis does decline with age, more so in females. Body weights of young males and females fed ketogenic diet for one week, along with middle-aged females increased by 1-1.5 grams after the 1-week diet, but body weights declined by 1.5 grams in both old male and female mice **(Extended Fig. 2c, d)**. Plasma BHB concentrations induced by ketogenic diet remained relatively consistent across all ages **(Fig. 3g)**, in contrast to both our prior fasting and control diet BHB blood concentration data. Insulin levels did not increase significantly with age on ketogenic diet **(Fig. 3h)**, and NEFA levels remained more stable as well **(Fig. 3i)**. We observed a dramatic drop in glucose levels in old males and females along with a respective increase in plasma BHB concentrations, exceeding a normal KD-range of ∼2.0 mM **(Fig 3j)**. Overall, these data suggest that aging mice retain the latent capacity to induce the ketogenic system under the strong stimulus of a ketogenic diet. In addition, similar trajectories were observed in males and females, leading us to investigate upstream regulators of ketogenesis next.

**Fig. 3.**
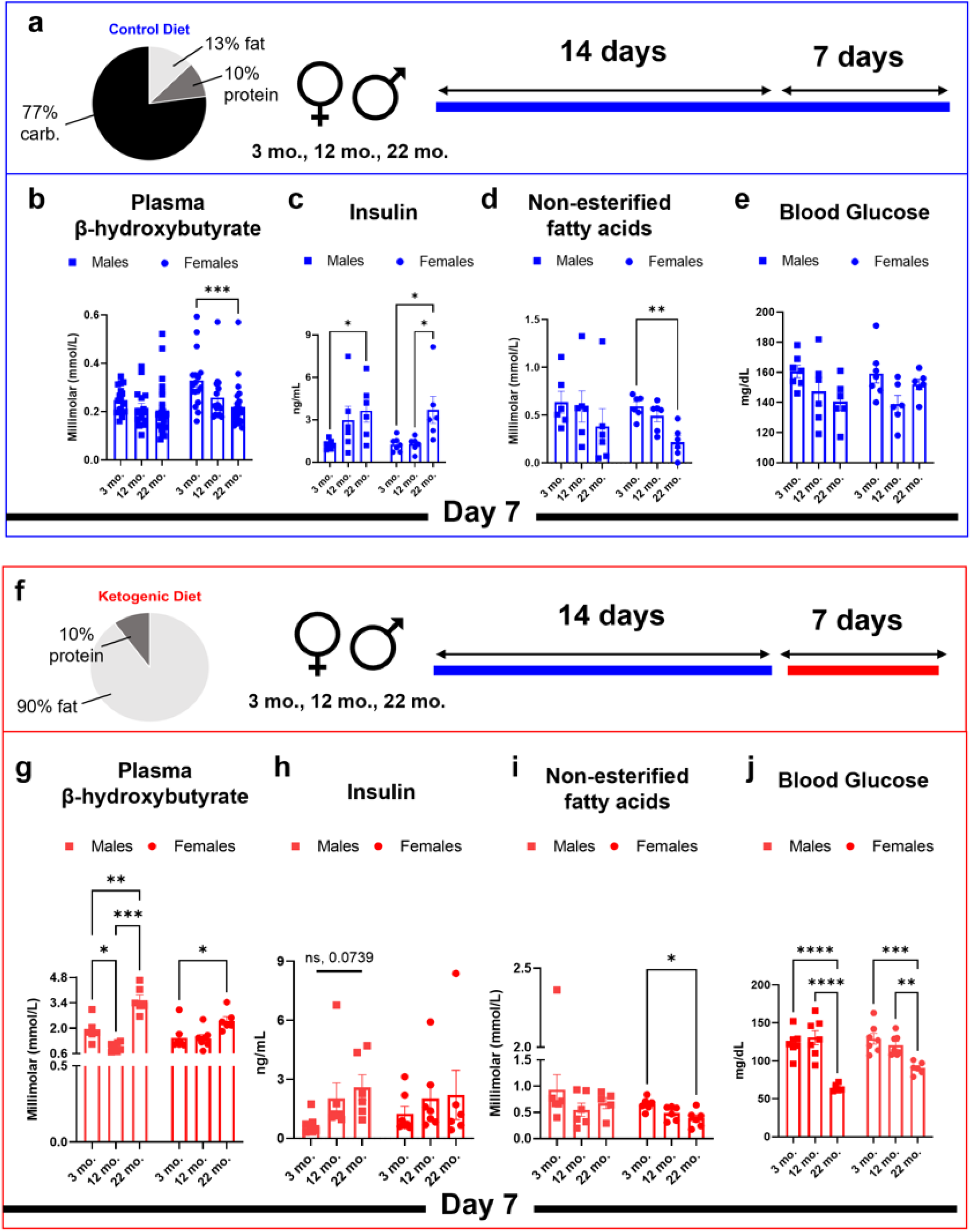
Ketogenic diet helps augment non-fasting ketosis in aging. **a**, Control diet, sex and age groups, and feeding timeline **b**, Day 7 plasma BHB concentrations of pooled cohort **c**, Day 7 insulin levels **d**, Day 7 non-esterified fatty acids levels **e**, Day 7 blood glucose levels **f**, Ketogenic diet, sex and age groups, and feeding timeline **g**, Day 7 plasma BHB concentrations **h**, Day 7 insulin levels **i**, Day 7 non-esterified fatty acids levels **j**, Day 7 blood glucose levels **b**, n=16-26 per sex groups **c-e, g-j**, n=6-7 per sex, diet and age groups, all data are presented as mean ± SEM; Ordinary two-way ANOVA, post-hoc Tukey’s for multiple comparisons test between age groups, *p<0.0332, **p<0.0021, ***p<0.0002. ****p<0.001.

### KD activates overall ketogenic capacity via upstream regulators of ketogenesis but declines with aging

Ketogenesis is tightly regulated by fatty acid availability and, transcriptionally, by the key regulator of fatty acid and ketone body metabolism, peroxisome proliferator activated receptor alpha (PPARα). To better understand how transcriptional regulation of ketogenesis changes with age and differs between sexes, we examined the gene expression of enzymes, transporters, and upstream regulators involved in ketogenesis in the liver via RT-qPCR. In control-fed males, we observed strong age-related declines in gene expression of several regulators of ketogenesis including PPARα, cAMP responsive element binding protein 3 like 3 (*Creb3l3*), the PPARα transcriptional target murine cytochrome P450, family 4, subfamily a, polypeptide 10 (*Cyp4a10*), phosphoenolpyruvate carboxykinase 1 (*Pck1)*, a target of gluconeogenesis and importantly, 3-Hydroxy-3-Methyl-CoA synthase 2, (*Hmgcs2)*, the rate-limiting enzyme of ketogenesis **(Fig. 4a, Extended Fig. 3a-n)**. In our higher-powered repeat control cohort, we were able to confirm the key age-related expression declines in *PPARα and Hmgcs2* **(Extended Fig. 4a-h)**. KD strongly induces ketogenic gene expression in males, but for several key genes the strength of induction declines with age, including *PPARα, Creb3l3, Cyp4a10*, murine cytochrome P450, family 4, subfamily a, polypeptide 14 (*Cyp4a14)*, and the ketogenesis pathways enzymes acetyl-CoA acetyltransferase (*Acat1*), 3-hydroxymethyl-3-methylglutaryl-CoA lyase (*Hmgcl*), and *Hmgcs2* **(Fig. 4a, Extended Fig. 3a-n)**. Interestingly, KD suppression of genes involved in fatty acid synthesis (a key feature of KD) such as fatty acid synthase (*Fasn*), acetyl-CoA carboxylase (*Acaa*), stearoyl-CoA desaturase (*Scd*), and ATP citrate lyase (*Acly*), was strong at all ages **(Fig. 4a)**. Mass spectrometric analysis (via data-independent acquisitions)^33,34^ comparing relative protein abundance between young KD vs young CD and old KD vs old CD, respectively, in the same liver samples revealed similar patterns at the protein level. We observed an increase in CYP4A10, CYP4A14, PCK1, ACAT1, HMGCS2, BDH1, and SLC16A1 in young and old KD-fed males and a decrease in FASN and ACYL **(Fig. 4c)**. In control-fed females, we saw weak age-related expression patterns for ketogenic genes and relatively less overall induction with KD **(Fig. 4b, Extended Fig. 5a-n)**. Several upstream regulators and ketogenic pathway enzymes showed peak control diet expression at middle age, but only *PPARα* and *Bdh1* retained this pattern in the replication cohort (**Extended Figure 6a-h)**. There was no consistent age-related pattern of KD gene expression induction, and less KD induction overall at any age other than for males. Similarly to males, we saw strong downregulation in genes related to fatty acid synthesis by KD across all ages **(Fig. 4b)**.

**Fig. 4.**
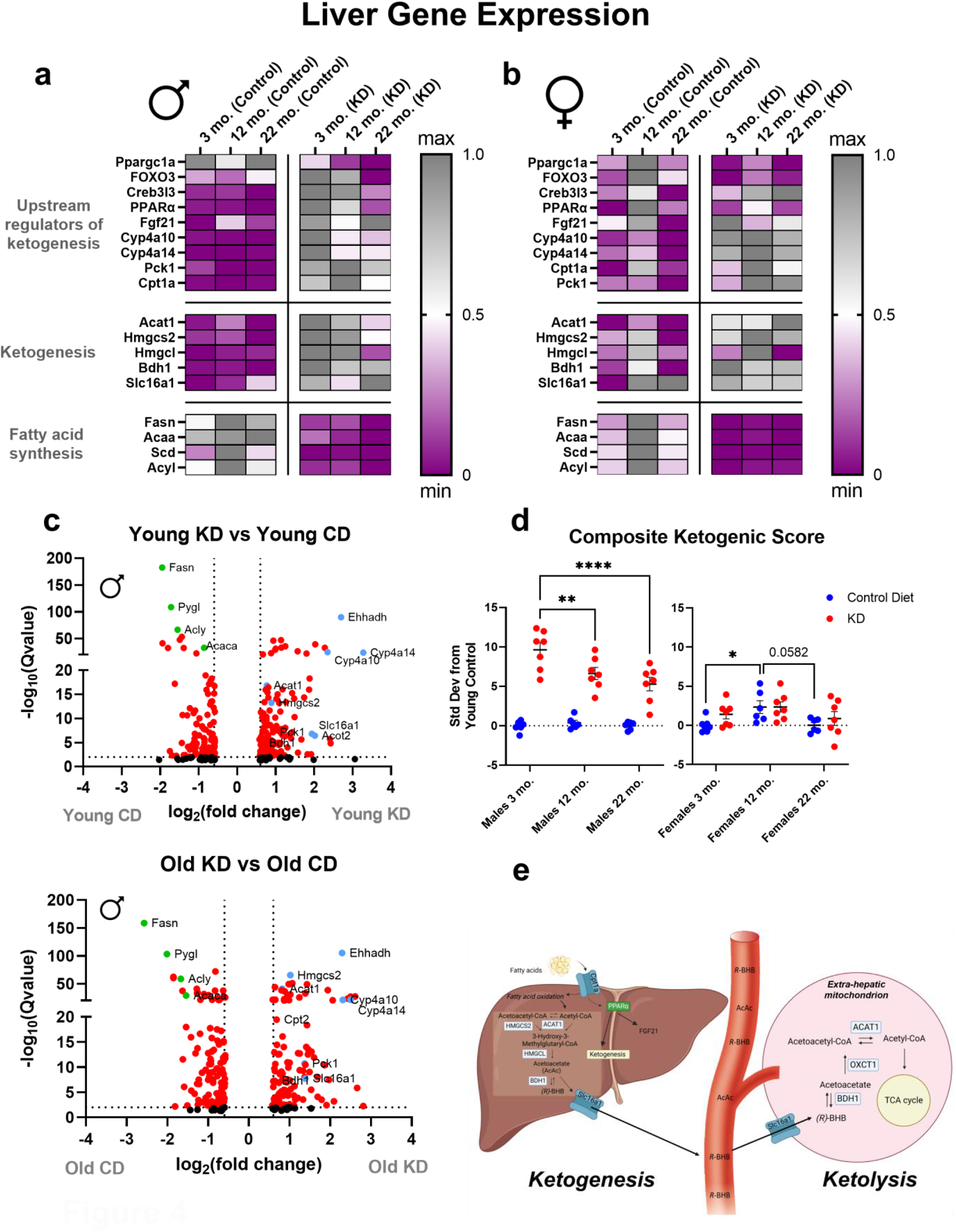
KD activates overall ketogenic capacity via upstream regulators of ketogenesis but declines with aging. **a**, Hepatic differential gene expression in control and KD-fed males **b**, Hepatic differential gene expression in control and KD-fed females **c**, Volcano plot of mass spectrometric analysis (via data-independent acquisitions) comparing relative protein abundance between young KD vs young CD and old KD vs old CD males (n=4/age and diet groups) **d**, Composite scores of control and KD-fed mice normalized to young mice **e**, schematic of regulators ketogenesis in the liver and ketolysis in extra-hepatic tissues **a, b**, **d**, n=6-7 per sex and age groups, all data are presented as mean ± SEM; Ordinary two-way ANOVA, post-hoc Tukey’s for multiple comparisons test between age groups, *p<0.0332, **p<0.0021, ***p<0.0002. ****p<0.001.

For a more systematic interpretation of these gene expression changes, we calculated a normalized composite score incorporating all of the upstream regulator and ketogenesis genes for each individual mouse **(Fig. 4d)**. Overall, ketogenic gene expression was relatively consistent across ages for control-fed males. KD strongly induced gene expression at all ages in males, but most strongly in young males. By contrast, females showed a life course trajectory of modestly higher ketogenic gene expression at middle age then lower at old age, but with little induction by KD at any age. Overall, our results show that males are more sensitive to gene expression changes by KD and that young male mice in particular are able to most strongly boost their ketogenic capacity through upregulation of PPARα and its targets compared to females and older males, similar to what we showed in our long term feeding of KD in old male mice (**Fig. 4e**)^20^.

### Tissue-specific changes in peripheral ketogenesis and ketolytic genes with age

Based on the changes we observed in hepatic ketogenesis, we investigated how these changes affect ketolysis in peripheral tissues like the heart and brain. In the heart, both control-fed males and females showed the highest expression of the rate-limiting ketolytic gene, 3-oxoacid CoA-transferase (*Oxct1*) along with the membrane transporter, solute carrier family member 1 (*Slc16a1*), at the oldest age **(Fig. 5a-b and Extended Fig. 7a-j)**. Interestingly, *Oxct1* was relatively suppressed by KD in the heart, while the key ketogenic gene *Hmgcs2* was strongly induced by KD in the oldest males and females **(Fig. 5a-b and Extended Fig. 7a-j)**. Overall, this suggests a pattern of compensatory increase in ketone body utilization with aging in the heart, along with possible augmentation by local ketogenesis under the stimulus of KD. In whole brain, only male mice showed an increase in ketolytic genes with age solute carrier family 16 member 7,(*Slc16a7*), *Acat1*, and *Oxct1*), not females **(Fig. 5c-d, Extended Fig. 8a-l)**. Both male and female brains showed a similar pattern of strong KD gene expression induction only at the youngest age, with stronger induction in females. Interestingly, this KD gene expression induction in young mice included the ketogenic enzyme *Hmgcs2*. Using LC-MS, we measured (R)-BHB in whole brains and observed relatively consistent amounts across ages and sexes under control diet feeding **(Fig. 5e-f)**, suggesting that homeostatic mechanisms preserve brain BHB availability despite age-related changes in plasma BHB concentrations. KD dramatically increased brain (R)-BHB in all ages and both sexes, but least so in middle-aged males. A third organ, kidney, showed age-related decline in ketolytic enzymes (*Oxct1* and *Bdh1*) in males but not females **(Extended Fig. 9a-j)**, along with strong induction by KD of *Hmgcs2* at all ages in both males and females. Altogether, we observed organ-specific patterns in age-related changes in the ketogenic system that likely represent the adaptive role of ketone-based energy production specific to each organ. Surprisingly, we more consistently observed induction of *Hmgcs2* by KD than ketolytic genes, suggesting a possible role for local ketogenesis within these tissues either for signaling roles or to augment an age-related decline in hepatic ketogenesis.

**Fig. 5.**
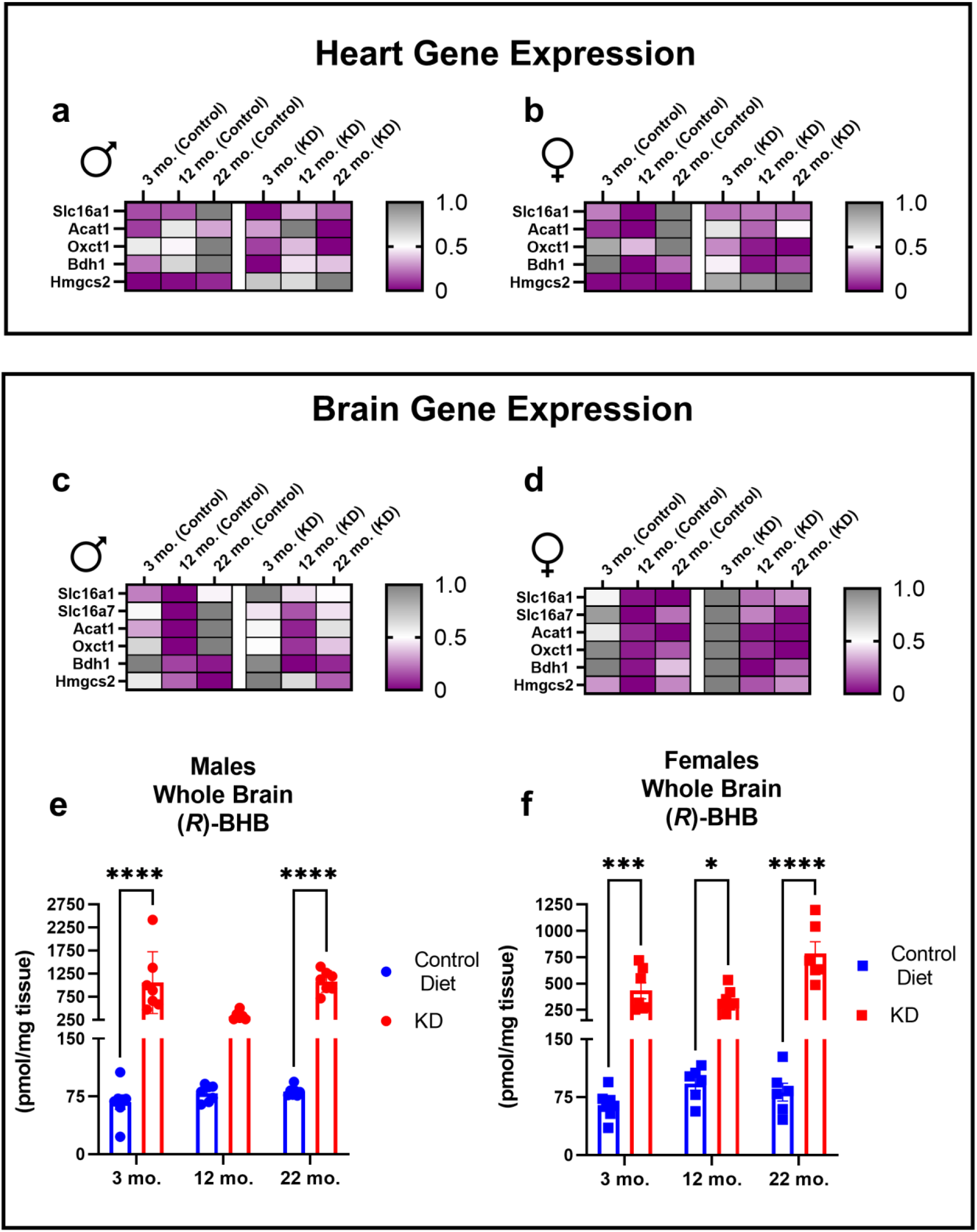
Tissue-specific changes in peripheral ketogenesis and ketolytic genes with age. **a**, Heatmap of heart gene expression in control and KD-fed males **b**, Heatmap of heart gene expression in control and KD-fed females **c**, Heatmap of brain gene expression in control and KD-fed males **d**, Heatmap of brain gene expression in control and KD-fed females **e**, Concentrations of *(R)*-BHB in whole brains of males **f**, Concentrations of *(R)*-BHB in whole brains of females **e-f**, n=6-7 per sex and age groups, all data are presented as mean ± SEM; Ordinary two-way ANOVA, post-hoc Tukey’s for multiple comparisons test between age groups, *p<0.0332, **p<0.0021, ***p<0.0002. ****p<0.001.

**Fig. 6.**
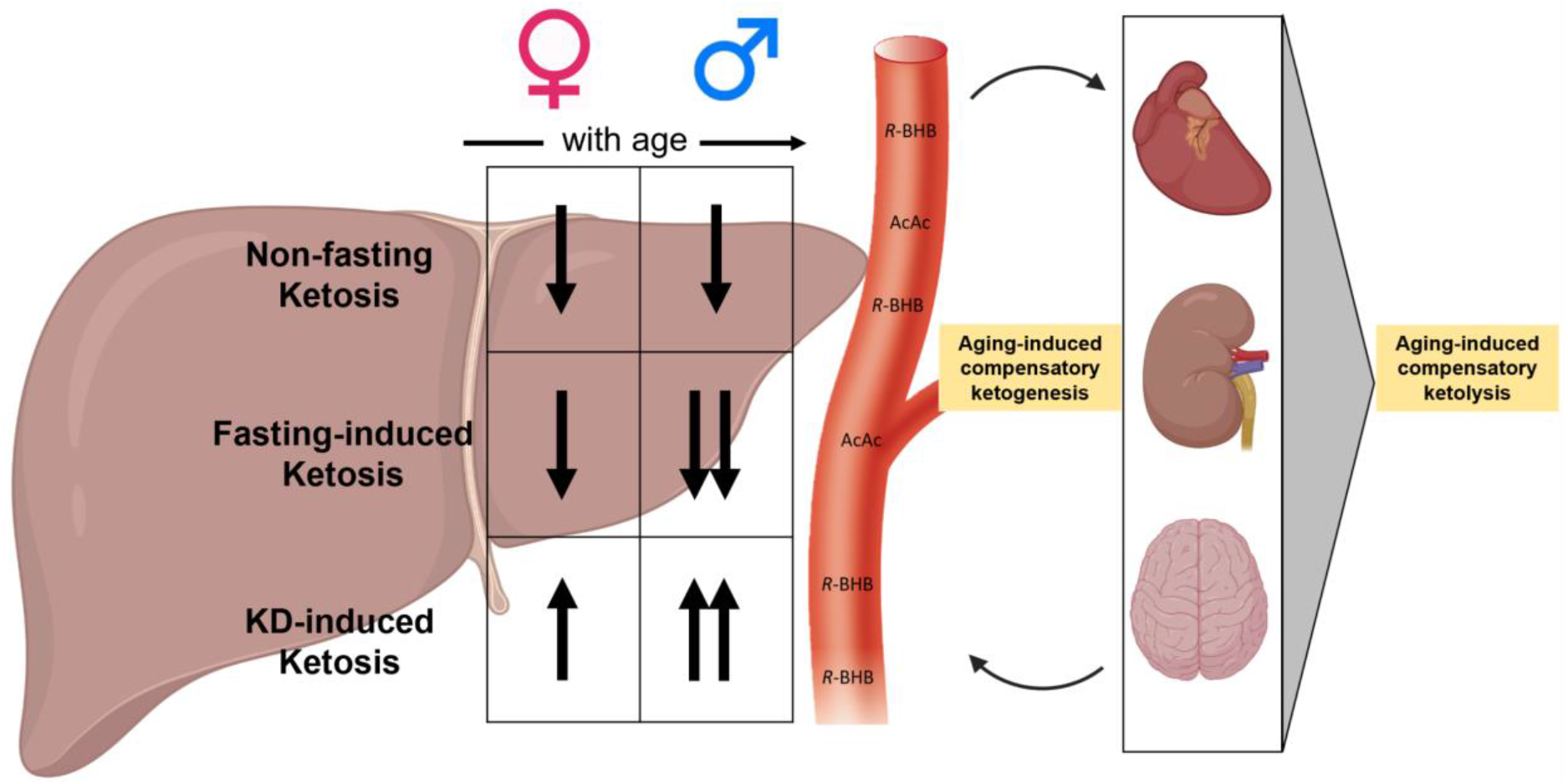
KD boosts the age-related declines non-fasting and fasting-induced ketosis in a sex-dependent manner. Non-fasted and fasted mice all showed a decline in circulating blood BHB with age, which was augmented by KD. The increases in blood BHB by KD were observed in both sex but was stronger in males. Changes in ketone body metabolism in extra-hepatic organs like the heart, kidney, and brain were diet-dependent. In control-fed animals, ketolysis was favored over ketogenesis whereas KD provided fats for tissue-specific ketogenesis. Overall, our work shows that hepatic ketogenesis declines in the liver with age, incapable of compensating for the hypometabolism of glucose we commonly see in aging. However, with KD, ketogenesis is promoted systemically to help compensate for this decline.

Finally, to directly test if induced extra-hepatic ketogenesis meaningfully contributes to blood BHB concentrations under KD, we generated a liver-specific knockout (KO) mouse of *Bdh1* (Albumin-Cre; *Bdh1*^fl/fl^) that should produce only AcAc, not BHB, from the liver^35^ **(Extended Fig. 10a)**. Unlike heart-specific *Bdh1* KO mice^36^ or whole-body *Bdh1* KO mice^37^, peripheral utilization of BHB should be unaffected in the mouse model. Body weights and blood glucose in these KOs should also remain unaffected **(Extended Fig. 10b-c, i-j)**. Ketogenesis, or perhaps only interconversion of AcAc to BHB, has been described in the kidney but under fasting conditions does not contribute to circulating BHB^38^. Surprisingly, we observed about 1/3 of normal blood BHB concentrations under KD in the liver KO **(Extended Fig. 10d**,**k)**, suggesting a role for non-hepatic ketogenesis under the strong stimulus of KD in compensating for loss of hepatic BHB production^35,39–41^. We confirmed efficacy of the knockout via RT-qPCR showing increased induction of *Hmgcs2* with KD, emphasizing ketogenesis, but an insignificant change in *Bdh1*, supporting the lack of AcAc to BHB conversion in livers of both male and female liver BDH1 KOs **(Extended Fig. 10e-h, l-o)**.

## Discussion

Studies showing lifespan extension through interventions like fasting, dietary and caloric restriction consistently show that metabolism and aging are tightly connected. Though these interventions hold promise in preserving healthy longevity, the role of pathways invoked by these interventions in normal aging often remains unclear. For an example, elevating the concentration of circulating ketone bodies in the blood via caloric restriction or ketogenic diet is associated with increased healthy longevity in mice, but whether ketone bodies change in aging under normal conditions has been understudied^16,42–44^. In this study, we systematically investigated ketosis in aging in C57BL/6N male and female mice under three conditions, a 16-hour fast, a typical high-carbohydrate diet and high-fat KD to recapitulate non-fasting and KD-induced ketosis respectively.

We observed three important sex differences that should critically inform mouse (and human) studies of metabolism in aging. First, during a 16-hour fast we observed higher concentrations of circulating plasma BHB in females than males, although both declined with age. Second, female RER is generally lower (more fat metabolizing) at baseline. Third, we only observed substantial gene expression induction by KD in males, despite similar non-fasting ketogenic responses. Our data are consistent with limited studies of sex differences in ketosis in humans. One clinical study showed fasting and postprandial plasma triacylglycerol concentrations were significantly higher in men, but women had higher plasma BHB^45^. Women have higher lipolytic release than men, which may help explain sexually dimorphic differences in lipid and glucose metabolism^4^. Several studies also revealed higher lipogenesis and lipid storage in females, hypothesized to be for reproduction, which could explain why female RER is lower^46,47^. A caveat to comparing mouse to human fasting studies is that mouse BHB concentrations exceeded 1.0 mM within 16 hours while heathy human adults reach that point only after 48-72 hours of fasting, likely due to the faster metabolic rate of mice versus humans^15,17,48^. Most metabolic studies in mice are undertaken only in young males, and our data show that the fasting/ketogenic response of young male mice is not representative of all ages and sexes. The relative lack of gene expression induction by KD in female mice was striking from a mechanistic standpoint. Both non-fasting and mechanistic studies in mice should include both sexes and, if targeting an age-related disease, use older mice whenever possible.

Our data also provide important clues to the heretofore obscure role of ketosis in aging. We hypothesized that ketogenic capacity might increase with age, in part due to reports of a compensatory role of ketone bodies in specific age-related diseases such as heart failure and Alzheimer’s disease. Mechanistic studies in mice have demonstrated that the failing heart adaptively switches to favor ketone body metabolism, and that provision of ketone bodies either exogenously or via KD improves heart function in heart failure models^25,36,49^. Instead we observed an age-related decline in BHB induction under fasting, a decline in non-fasting blood BHB concentrations, and a decline in expression of many ketogenic genes in the liver. Our data are consistent with a more limited but cross-sectional aging study in male mice that also showed that glucose and BHB were lower in the serum in the old, suggesting reduced production from the liver^50^. Adipose tissue is a critical organ for nutrient sensing and energy storage and has been widely studied to explain changes in metabolic health. Rising insulin levels and declining circulating NEFA possibly through decreased lipolysis in aging may proximately explain declining blood BHB levels with age^51–53^. We also suspect that the high content of carbohydrates in our control diet and the rising insulin levels with age could also inhibit ketogenic capacity through mTOR activation^44,54^. Aging also induces inflammatory activation of NLRP3 and NF-κB, releasing pro-inflammatory markers like IL-6 and TNFα, which could also diminish compensatory ketogenesis through PPARα inhibition^55^. Despite the many KD studies done mainly in males (mice and humans), a study administering exogenous ketone supplements in rats showed that age and sex both modulate circulating blood BHB. In older rats, the ketone ester increased concentrations of blood BHB more efficiently in male rats, supporting our data for the KD-fed males^56^.

In our study, we did find gene expression evidence of age-related compensatory utilization of ketone bodies in extrahepatic organs, despite the overall age-related decline in hepatic ketogenesis. In age-related diseases like heart failure and Alzheimer’s Disease, tissue-specific ketone body utilization increases, and overall utilization is determined by the circulating concentrations of ketone bodies in the blood ^36,39,57–61^. From our data, heart and brain, but not kidney, showed increased expression of ketolytic genes with age, but more so in control-fed animals, underscoring the importance of ketone body utilization in heart failure and the aging brain. Despite the levels being measured in whole brain, higher expression levels of the astrocyte specific monocarboxylate transporter, *Slc16a1 and* neuron-specific monocarboxylate transporter, *Slc16a2* also support an age-related increase in brain ketolysis^62–64^.

However, when given a strong stimuli like KD, ketogenesis through *Hmgcs2* gene expression dramatically increases in heart and kidney across all ages, suggesting a role for compensatory tissue-specific ketogenesis. More specifically, heart fibroblast and kidney mesangial cells have been shown to express *Hmgcs2*^65,66^. Cortex glia in *Drosophila melanogaster* have been shown to also participate in ketogenesis through AMPK signaling^67^. Based on our measurements of brain R-BHB, we suspect that the higher concentrations could help explain higher ketogenic activity to meet brain energetic demands^68–71^. Our previous study also showed that old male mice fed the same KD used in this study for 12 months had improved cognitive function, which was absent in mice fed both the high-carbohydrate and high-fat diets^21,72^. Further, we found that knocking out liver *Bdh1*, which prevents liver production of BHB, still left ∼1/3 of circulating BHB intact under KD, suggesting that extrahepatic ketogenesis under KD is a meaningful contributor to total body BHB production. In addition to providing extra-hepatic tissues with an alternative energy substrate, ketone bodies also have important signaling properties. One paper showed that AcAc can protect the liver from high-fat-diet-induced fibrosis through mediating metabolic plasticity in macrophages^73^. A study also showed that ketone bodies have an important role in the gut, such as mediating intestinal stem cell homeostasis under different responses to diets^74^.

This study is the first to systematically examine ketone body metabolism across ages and sexes. The power of our cohorts limited the detection of metabolic and gene expression changes to some extent, and so key measures were validated in a repeat, larger, non-fasting control-fed cohort. Although we were able to distinguish between ketogenesis and ketolysis through gene expression analyses, ketone production and oxidation rates are not determined solely by gene expression. Future experiments directly measuring oxidation rates using stable isotope labeling across sexes and ages will deepen our understanding of the changes in ketone body metabolism. We also acknowledge that in mice, sexes, ages, and strains can all influence the effects of dietary interventions^75,76^. While KD remains to be an effective therapeutic concept in brain aging, the driver of the sexual dimorphic changes we observed in our study requires further elucidation. We also have reasons to believe that sex hormones, which were not accounted for, regulate hepatic transport of ketone bodies and could help explain the sexual dimorphic changes in ketone body metabolism^77^. Overall, this study identifies critical age- and sex-differences in ketone body metabolism under fasting, non-fasting, and ketogenic diet conditions, and suggests the potential of interventions that target the endogenous ketogenic system to treat age-related diseases.

## Methods

### Mouse strains, housing and husbandry

C57BL/6 mice were obtained from the National Institute on Aging’s Aged Rodent Colony. *Bdh1*^fl/fl^ mice were previously published^36^. Albumin-Cre mice^78^ were obtained from the Jackson Laboratory (#003574). All mice were maintained in animal facility under a 6:00 am to 6:00 pm light cycle. All mice were housed in groups before being single caged for their respective diets for one week. All mice were maintained according to the National Institutes of Health guidelines and all procedures and protocols were approved by the Institutional Animal Care and Use Committee at Buck Institute for Research on Aging (Novato, CA).

### Mouse diets and feeding

Food was provided ad libitum at all times. Per-calorie macronutrient content for customized diets (Envigo) is as follows: Control, 10% protein, 13% fat, and 77% carbohydrate (TD.150345); KD 10% protein and 90% fat (TD.160153). Both diets are matched on a per-calorie basis for micronutrient content, fiber, and preservatives. All mice were acclimated on a control diet for 2 weeks in the groups they arrived before they were single caged for this study.

### Metabolic Cage

C57BL/6 male and female mice (3, 12, 22 months) were single-caged in metabolic chambers (Promethion Cages from Sable Systems International) for 108 hours. Raw data files were analyzed using CalR (https://CalRapp.org).

### Plasma Beta-Hydroxybutyrate Measurements

Blood was obtained via distal tail-snip (10-40 uL) or cardiac puncture if the animal was euthanized for tissue collection in lithium-heparin coated microvettes (Sarstedt CB 300 LH). Blood was spun at 1500 x g for 15 mi at 4C and kept in -80C until thawed for assay. Plasma BHB levels were measured using StanBio LiquiColor Beta-Hydroxybutyrate colorimetric test per manufacturer’s protocol (2440-058).

### Glucose Measurements

Glucose levels were measured using a Accu-Chek blood glucose meter. Glucose measurements for Figure 1 was every 2 hours during the 16-hour fast (6am-10pm). Glucose measurements for Figure 3 was done before light period (6am) following mice euthanasia.

### Insulin Measurements

Insulin levels were measured using Mercodia Ultrasensitive Mouse Insulin ELISA per manufacturer’s protocol (Mercodia Cat. 10-1249-01).

### Non-esterified fatty acid (NEFA) Measurements

NEFA levels were measured using NEFA Randox ELISA per manufacturer’s protocol (Randox Cat. FA115).

### Real-time quantitative PCR

Mice were euthanized before the light period (6am) at the end of the 1-week feeding period and tissues were immediately collected and snap-frozen for -80°C storage. Snap frozen tissues were homogenized using RiboZol RNA Extraction Reagent (VWR; N580) and Next Advance Bullet Blender (BBY24M). RNA was extracted using the Qiagen RNEasy Mini Kit (Qiagen Cat. 169012183) and checked for purity using Thermo Scientific Nanodrop 2000 (Serial No. D631). cDNA synthesis was carried out with Superscript cDNA Synthesis Kit (BioRad Cat. 1708891) and real-time quantitative PCR was performed using iTaq Universal SYBR Green Supermix (BioRad Cat. 1725121) in BioRad CFX96 Real-Time System. Gene expression analyses were normalized to either *B2m* (liver) or *Rplp0* (heart, brain, kidney) housekeeping genes.

### Absolute Quantification of brain (R)-BHB

Frozen whole brains (∼30-50 mg) were homogenized in extraction buffer [3:1, v/v acetonitrile and HPLC-grade H2O] with Next Advance Bullet Blender (BBY24M). Derivatization was adapted from the following paper^79^. Extracted samples were dried using DNA SpeedVac System (ThermoFisher Scientific Model DNA130-115) and resuspended in 98:2 H2O:Methanol containing .1% formic acid, mixed and centrifuged at 10,000*xg* for 10 minutes. Supernatant was then transferred into HPLC vials. A sample volume of 2 μL was injected into the UPLC-MS/MS Thermo Q Exactive with Vanquish Horizon in Full MS and PRM scan modes using positive ionization. The analysis was performed on a Accucore Vanquish C18+ column (100 × 2.1 mm, 1.5μm particle size; ThermoFisher Cat. #20073385). The following mobile phases were used: A) HPLC-graded H2O containing 0.1% (v/v) formic acid and B) methanol containing 0.1% (v/v) formic acid with the gradient starting at 2% B for 0.5 min and gradually increasing to 10.4%B until 10.0 min, 2%B until 10.1 min, and 2%B until 12.0 at 0.150 mL/min flow rate for a total run time of 12.0 minutes. Column was maintained at 40°C. LC system was hyphenated to Thermo Q Exactive MS equipped with heated electrospray ionization (HESI) source. The MS system was operated in Full Scan MS or PRM modes using positive ionization. MS scan range was 50.0 to 750.0 m/z in Full MS scan mode. The resolution was set to 140,000 with AGC target 3e6, isolation window 1.0 m/z and optimal collision energy was 50 (arbitrary units). For PRM scan mode the isolation window was set to 0.4 m/z and resolution was set to 70,000 with AGC target 1e6. Common HESI parameters were auxiliary gas: 5, sheath gas flow: 50, sweep gas: 0, spray voltage 3 kV, capillary temperature 320°, S-lens 55.0, and auxiliary gas temperature: 150°C. Quantification of area response ratios were processed and acquired using Thermo Scientific Xcalibur software (OPTON-30965). Area Response Ratios (ES/IS) from three technical replicates were averaged, and a simple regression line was constructed. To quantify the ‘samples’, Area Response Ratios (ES/IS) from three technical replicates were averaged and concentrations were calculated using the Prism-generated calibration curve equations. Amounts were calculated as [(100*concentration) pmol]/[mg tissue weight].

### Proteomics Analysis, Protein Digestion and Desalting

Livers of wild type and mice fed a ketogenic diet both young and old, with 4 biological replicates each, were homogenized in 800 μL of lysis buffer containing 8 M urea, 200 mM triethylammonium bicarbonate (TEAB), pH 8, 75 mM sodium chloride, 1 μM trichostatin A, 3 mM nicotinamide, and 1x protease/phosphatase inhibitor cocktail (Thermo Fisher Scientific, Waltham, MA), with 2 cycles in a Bead Beater TissueLyser II (Qiagen, Germantown, MD) at 24 Hz for 2 min each. Lysates were clarified by spinning at 15,700 x g for 15 min at 4°C, and the supernatant containing the soluble proteins was collected. Protein concentrations were determined using a Bicinchoninic Acid Protein (BCA) Assay (Thermo Fisher Scientific, Waltham, MA). Protein lysates (3.1 mg each) were reduced using 20 mM dithiothreitol (DTT) in 50 mM TEAB for 30 min at 37 °C, and after cooling to room temperature, alkylated with 40 mM iodoacetamide (IAA) for 30 min at room temperature in the dark. Samples were diluted 4-fold with 50 mM TEAB, pH 8, and proteins were digested overnight with a solution of sequencing-grade trypsin (Promega, San Luis Obispo, CA) in 50 mM TEAB at a 1:50 (wt:wt) enzyme:protein ratio at 37°C. This reaction was quenched with 1% formic acid (FA) and the sample was clarified by centrifugation at 2,000 x g for 10 min at room temperature. Clarified peptide samples were desalted with Oasis 30-mg Sorbent Cartridges (Waters, Milford, MA). 100 μg of each peptide elution were aliquoted for analysis of relative protein-level changes, after which all desalted samples were vacuum dried. The 100 μg whole protein lysate aliquots were re-suspended in 0.2% FA in water at a final concentration of 1 μg/μL. Finally, indexed retention time standard peptides (iRT; Biognosys, Schlieren, Switzerland)^80^ were spiked in the samples according to manufacturer’s instructions.

### Mass Spectrometric Acquisitions

Reverse-phase HPLC-MS/MS analyses were performed on a Dionex UltiMate 3000 system coupled online to an Orbitrap Exploris 480 mass spectrometer (Thermo Fisher Scientific, Bremen, Germany). The solvent system consisted of 2% ACN, 0.1% FA in water (solvent A) and 80% ACN, 0.1% FA in ACN (solvent B). Digested peptides (400 ng) were loaded onto an Acclaim PepMap 100 C_18_ trap column (0.1 × 20 mm, 5 μm particle size; Thermo Fisher Scientific) over 5 min at 5 μL/min with 100% solvent A. Peptides (800 ng) were eluted on an Acclaim PepMap 100 C_18_ analytical column (75 μm x 50 cm, 3 μm particle size; Thermo Fisher Scientific) at 300 nL/min using the following gradient: linear from 2.5% to 24.5% of solvent B in 125 min, linear from 24.5% to 39.2% of solvent B in 40 min, up to 98% of solvent B in 1 min, and back to 2.5% of solvent B in 1 min. The column was re-equilibrated for 30 min with 2.5% of solvent B, and the total gradient length was 210 min. Each sample was acquired in data-independent acquisition (DIA) mode^33,34^. Full MS spectra were collected at 120,000 resolution (Automatic Gain Control (AGC) target: 3e6 ions, maximum injection time: 60 ms, 350-1,650 *m/z*), and MS2 spectra at 30,000 resolution (AGC target: 3e6 ions, maximum injection time: Auto, Normalized ^33,34^Collision Energy (NCE): 30, fixed first mass 200 *m/z*). The isolation scheme consisted of 26 variable windows covering the 350-1,650 *m/z* range with an overlap of 1 *m/z*^81^.

### Data Analysis, DIA-MS Data Processing and Statistical Analysis

DIA data was processed in Spectronaut (version 16.0.220524.53000) using directDIA. Data was searched against a database containing all *Mus musculus* entries extracted from UniProtKB-TrEMBL (86,521 protein entries; August 24, 2021). Trypsin/P was set as digestion enzyme and two missed cleavages were allowed. Cysteine carbamidomethylation was set as fixed modification, and methionine oxidation, and protein N-terminus acetylation variable modifications. Data extraction parameters were set as dynamic. Identification was performed requiring a 1% q-value cutoff on the precursor ion and protein level. For the protein lysate samples, quantification was based on the peak areas of extracted ion chromatograms (XICs) of 3 – 6 MS2 fragment ions, and local normalization was applied. iRT profiling was selected. Quantification was based on the peak areas of XICs of 3 – 6 MS2 fragment ions, without normalization as well as data filtering using q-value and imputation of missing values with empirical noise.

Differential expression analysis comparing ketogenic diet to wild-type mouse livers was performed using a paired t-test, and p-values were corrected for multiple testing, specifically applying group wise testing corrections using the Storey method^82^. Quantifiable proteins consist of protein groups with at least two unique peptides and significantly-altered proteins have a q-value < 0.01 and absolute Log_2_(fold-change) > 0.58 (Supplementary Tables).

## Data Availability

Raw data and complete MS data sets have been uploaded to the Mass Spectrometry Interactive Virtual Environment (MassIVE) repository, developed by the Center for Computational Mass Spectrometry at the University of California San Diego, and can be downloaded using the following link: https://massive.ucsd.edu/ProteoSAFe/dataset.jsp?task=934f911545e84050ac0c18346d12006c (MassIVE ID number: MSV000090365; ProteomeXchange ID: PXD036921). Enter the username and password in the upper right corner of the page:

Username: *MSV000090365_reviewer*

Reviewer password: *winter*

### Heatmap Differential Expression Analysis

An average value was calculated for each gene across ages of both control and KD groups of each sex. Values are to represent minimum and maximum differential expressions for each gene. This analysis does not provide statistical significance, but rather allowed for a better representation of KD-induction compared to our control groups.

### Ketogenic Composite Score Analysis

Composite scores were calculated to represent an overall trajectory of ketogenic genes shown on the heatmap. The standard deviations and means of young control diet from each respective sex were normalized and used to calculate ketogenic scores for 12 and 22 months of both control and KD groups. Each individual mouse was normalized as the number of baseline standard deviations from the baseline mean.

### Quantification and Statistical analysis

Figures are presented as mean ± SEM, P values are calculated by either one-way ANOVA or two-way ANOVA with additional post-hoc testing for group comparisons. Statistical calculations are carried out in Prism 9.4.1 (GraphPad Software, San Diego, California USA, www.graphpad.com). Exact replicate numbers “N” are specified in the figure legends for each experiment, and refer to the number of individual mice.

## Acknowledgements

This work was supported by NIH grant R01AG67333 (J.C.N.), and Buck intramural funds (J.C.N.), NIA T32 AG052374 (B.E), NIA T32 AG000266 (M.N), and NIH R01HL151345 (D.P.K). We thank S. Galkina from Buck Institute’s Phenotyping Core for administering the metabolic cage study, A. Holcom for her assistance in the repeat control cohort tissue collection, and B. J. Stubbs for critical reading of the manuscript. Some figure illustrations were created with BioRender.com.

## Authorship contributions

B.E. conceived the studies, designed the experiments, analyzed experiments, and wrote the manuscript. M.N., O.P., and T.Y.G helped perform experiments. J.P.R., C.D.K., and B.S. performed and analyzed proteomics experiments. T.C.L. and D.P.K generated Bdh1^fl/fl^ mice. D.P.K provided important insights and suggestions to manuscript. J.C.N. supervised the study and co-wrote the manuscript.

## Additional Information

All data needed to evaluate the conclusions in the paper are present in the paper and Extended Figures. Data requests should be submitted to the corresponding author.

