## Extended Figures for "Ketone body metabolism declines with age in mice in a sex-dependent manner"

Eap B et al.

### Extended Figures 1-10

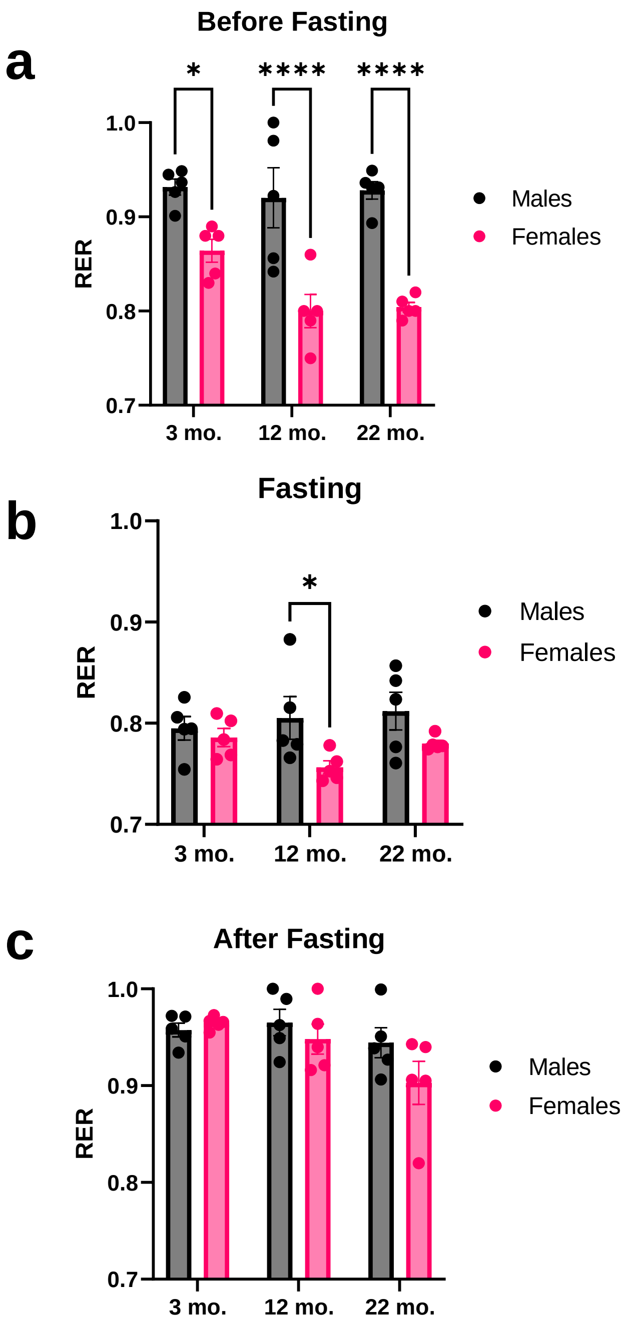


**Extended Data Fig. 1. Sex differences in metabolic state.**
**a,** RER values of males and females before fasting **b,** RER values of males and females during 24-hour fasting **c,** RER values of males and females after 24-hour fasting **a-c,** n=4-5 per age and sex groups, all data are presented as mean ± SEM; Ordinary two-way ANOVA, post-hoc for Šídák's multiple comparisons test between sex groups, *p<0.0332, **p<0.0021, ***p<0.0002. ****p<0.001.

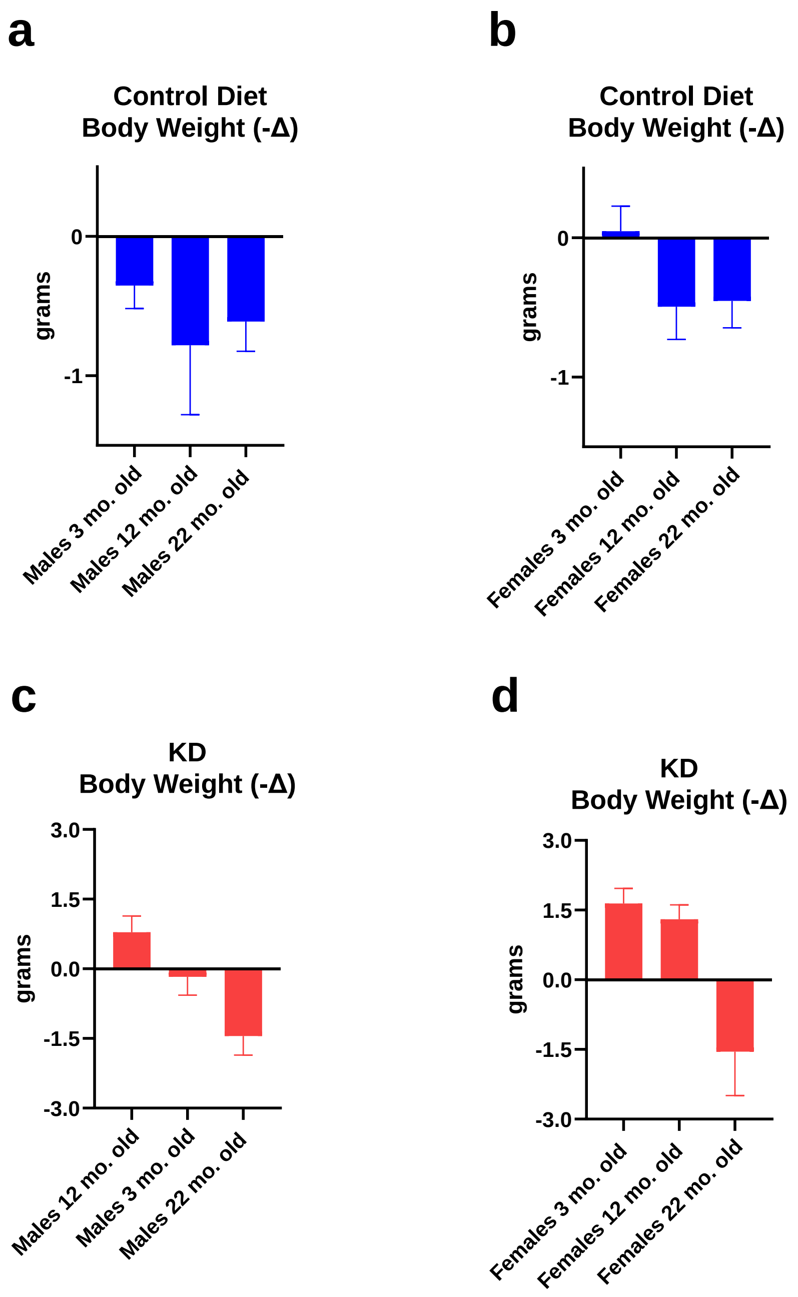


**Extended Data Fig. 2. Overall change in body weight after 1-week feeding.**
**a,** Body weight changes in control-fed males **b,** Body weight changes in control-fed females **c,** Body weight changes in KD-fed males **d,** Body weight changes in KD-fed females. **a-d,** All data are presented as mean ± SEM.

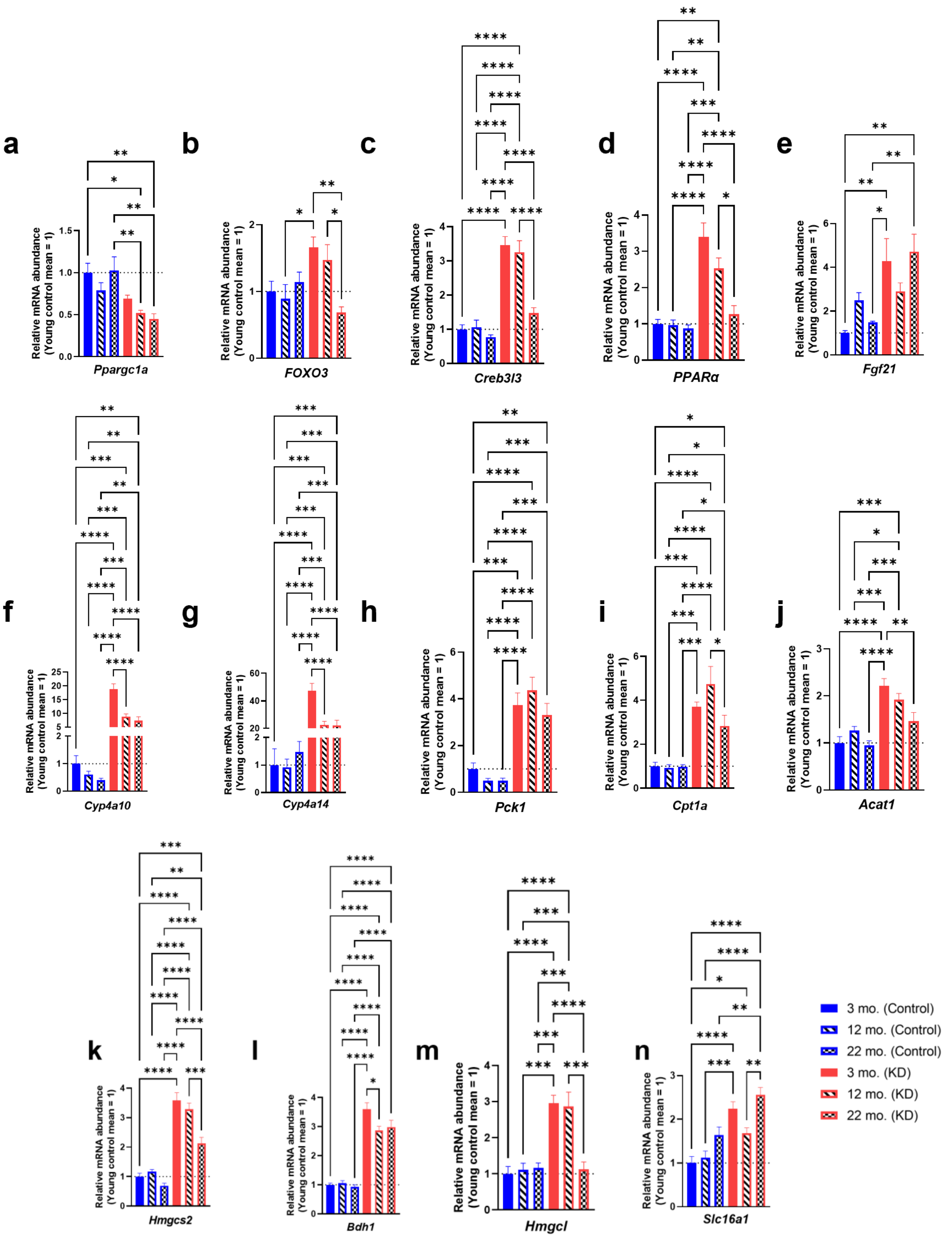


**Extended Data Fig. 3. Gene expression of upstream regulators of ketogenesis and ketogenesis in control and KD-fed male livers.**

**a-n,** *Ppargc1a, Foxo3, Creb3l3, Pparα, Fgf21, Cyp4a10, Cyp4a14, Pck1, Cpt1a, Acat1, Hmgcs2, Bdh1, Hmgcl, Slc16a1* gene expression of day 7 control and KD-fed male mice; n=6-7 per age and diet groups; all data are presented as mean ± SEM; Ordinary one-way ANOVA, Tukey's multiple comparisons test between age and diet groups; *p<0.0332, **p<0.0021, ***p<0.0002. ****p<0.001.

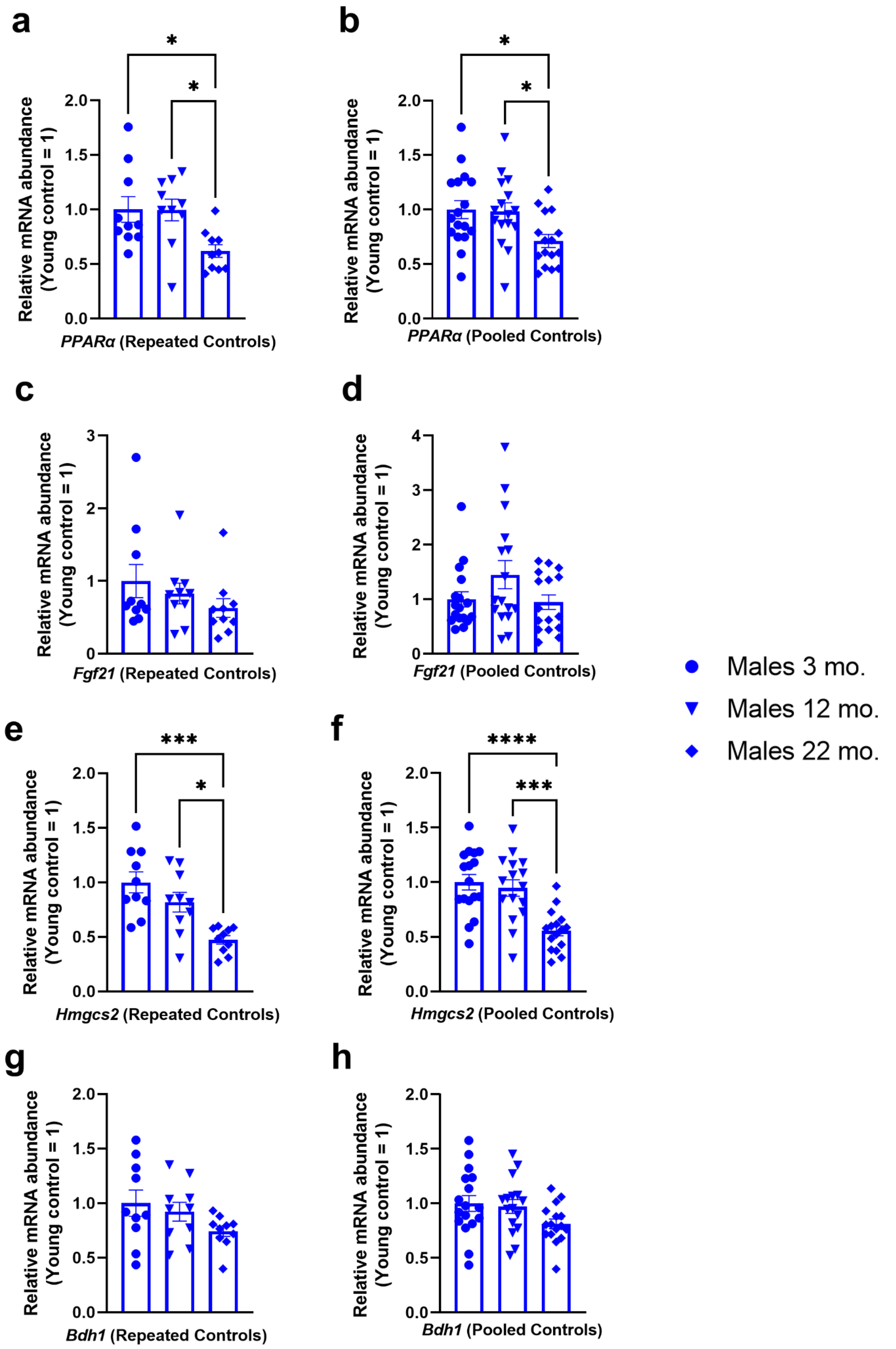


**Extended Data Fig. 4. Liver gene expression of control-fed males (repeated cohort) and pooled (original and repeated) cohorts.**
**a,c,e,g,** *Ppargc1a, Fgf21, Hmgcs2, Bdh1* gene expression of day 7 repeated control-fed males; n=10 per age group. **b,d,f,h,** *Ppargc1a, Fgf21, Hmgcs2, Bdh1* gene expression of day 7 pooled (pilot and repeated control-fed males; n=16-17 per age group. **a-h,** All data are presented as mean ± SEM; Ordinary one-way ANOVA, Tukey's multiple comparisons test between age groups; *p<0.0332, **p<0.0021, ***p<0.0002. ****p<0.001.

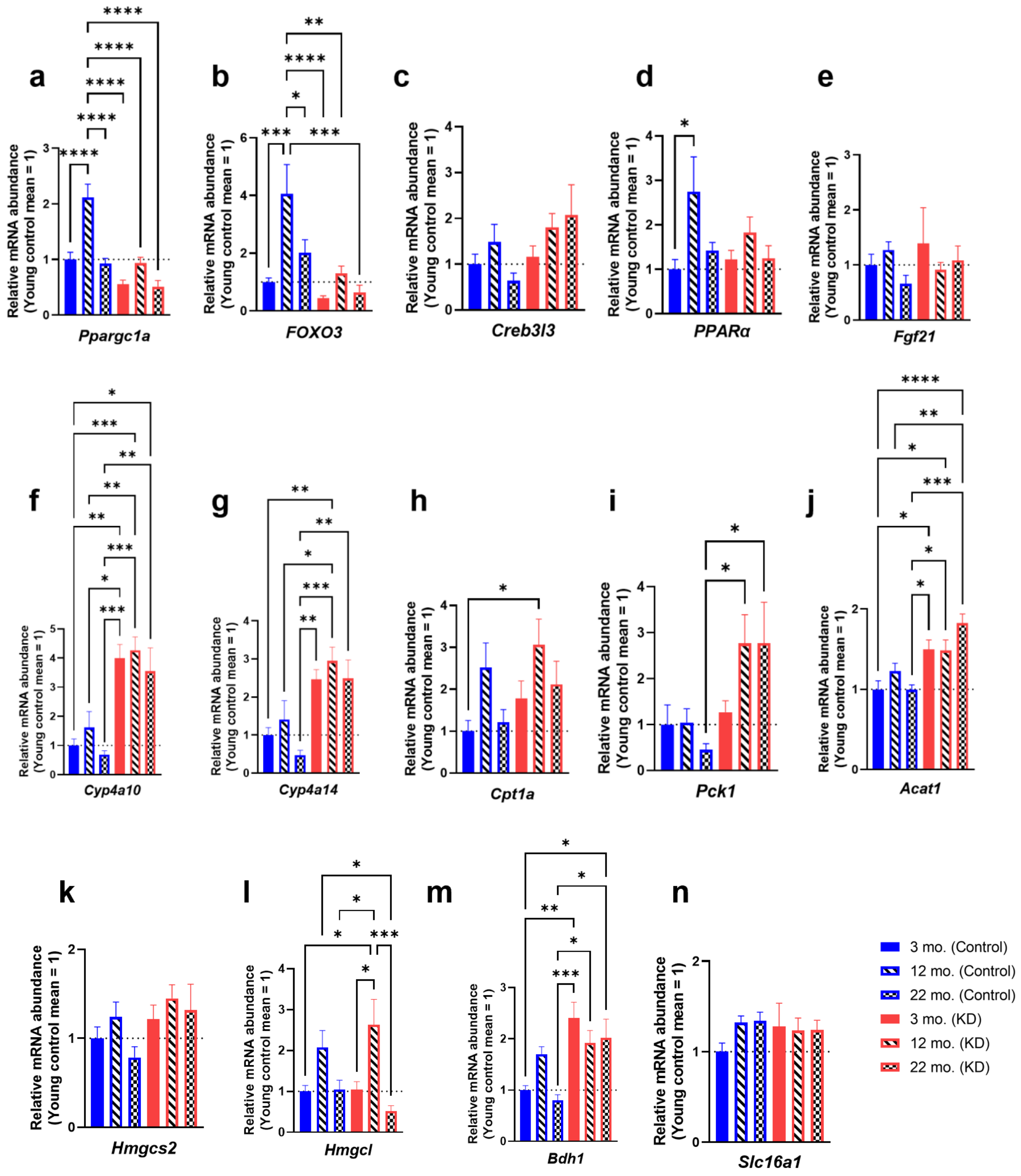


**Extended Data Fig. 5. Gene expression of upstream regulators of ketogenesis and ketogenesis in control and KD-fed female livers.**

**a-n,** *Ppargc1a, Foxo3, Creb3l3, Pparα, Fgf21, Cyp4a10, Cyp4a14, Cpt1a, Pck1, Acat1, Hmgcs2, Bdh1, Hmgcl, Slc16a1* gene expression of day 7 control and KD-fed female mice; n=6-7 per age and diet groups; all data are presented as mean ± SEM; Ordinary one-way ANOVA, Tukey's multiple comparisons test between age and diet groups; *p<0.0332, **p<0.0021, ***p<0.0002. ****p<0.001.

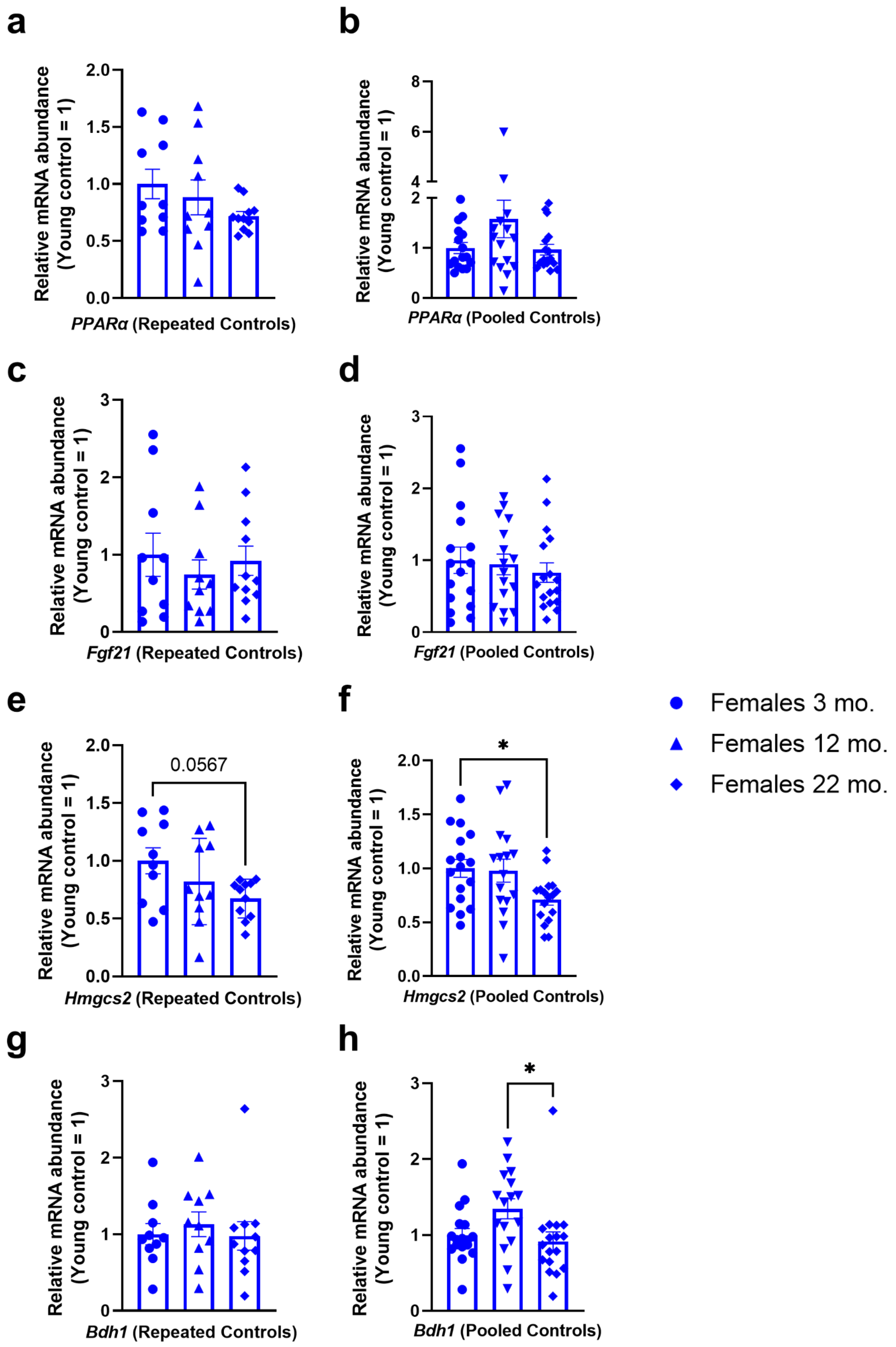


**Extended Data Fig. 6. Liver gene expression of control-fed females (repeated cohort) and pooled (original and repeated) cohorts.**

**a,c,e,g,** *Ppargc1a, Fgf21, Hmgcs2, Bdh1* gene expression of day 7 repeated control-fed females; n=10-11 per age group **b,d,f,h,** *Ppargc1a, Fgf21, Hmgcs2, Bdh1* gene expression of day 7 pooled (pilot and repeated control-fed females); n=16-17 per age group. **a-h,** All data are presented as mean ± SEM; Ordinary one-way ANOVA, Tukey's multiple comparisons test between age groups; *p<0.0332, **p<0.0021, ***p<0.0002. ****p<0.001.

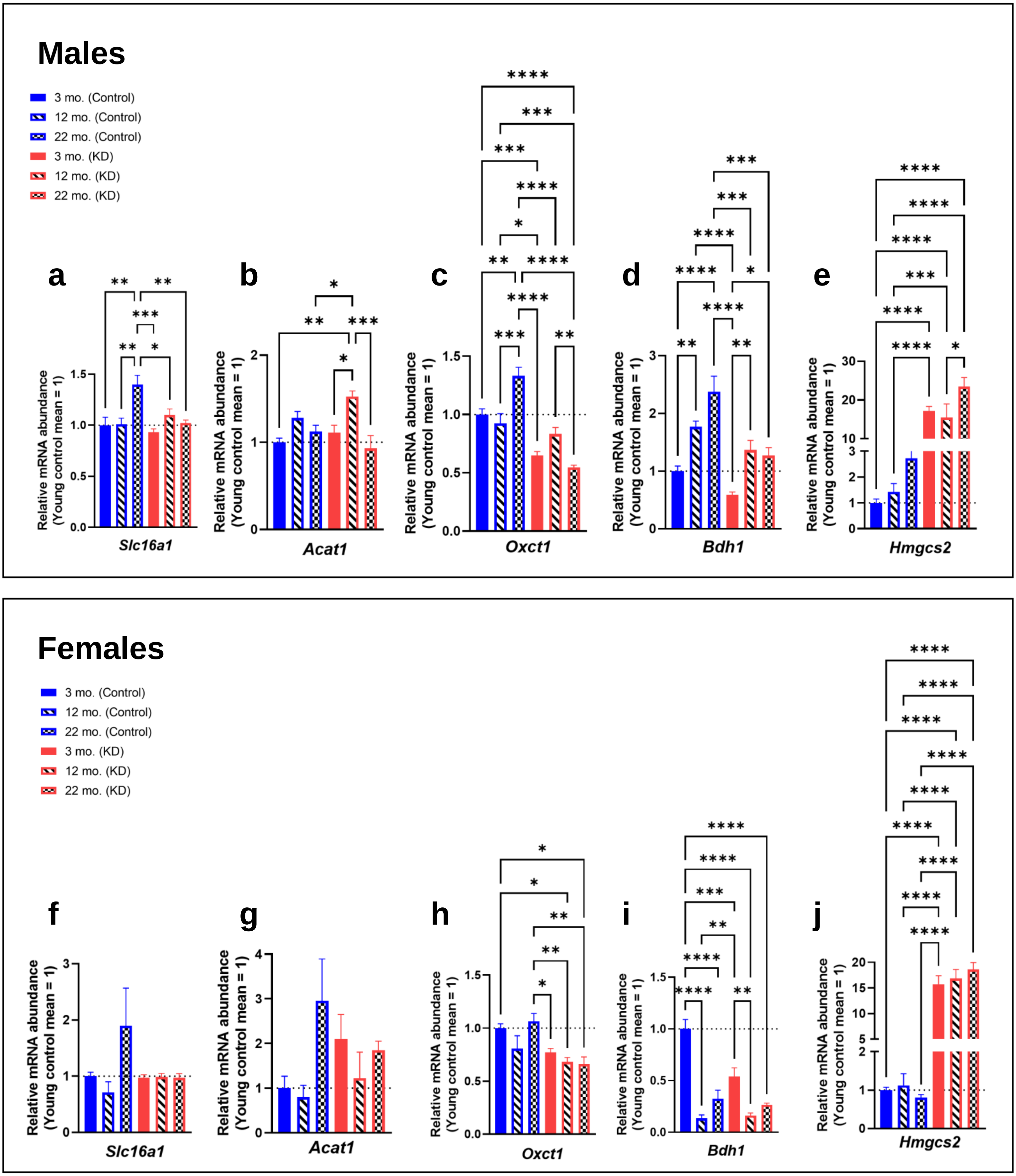


**Extended Data Fig. 7. Gene expression of ketone body metabolism genes in heart.**
**a-e,** *Slc16a1, Acat1, Oxct1, Bdh1, Hmgcs2* gene expression of day 7 control and KD-fed male mice; n=6-7 per age and diet groups **f-j,** *Slc16a1, Acat1, Oxct1, Bdh1, Hmgcs2* gene expression of day 7 control and KD-fed female mice; n=6-7 per age and diet groups **a-j,** All data are presented as mean ± SEM; Ordinary one-way ANOVA, Tukey's multiple comparisons test between age and diet groups; *p<0.0332, **p<0.0021, ***p<0.0002. ****p<0.001.

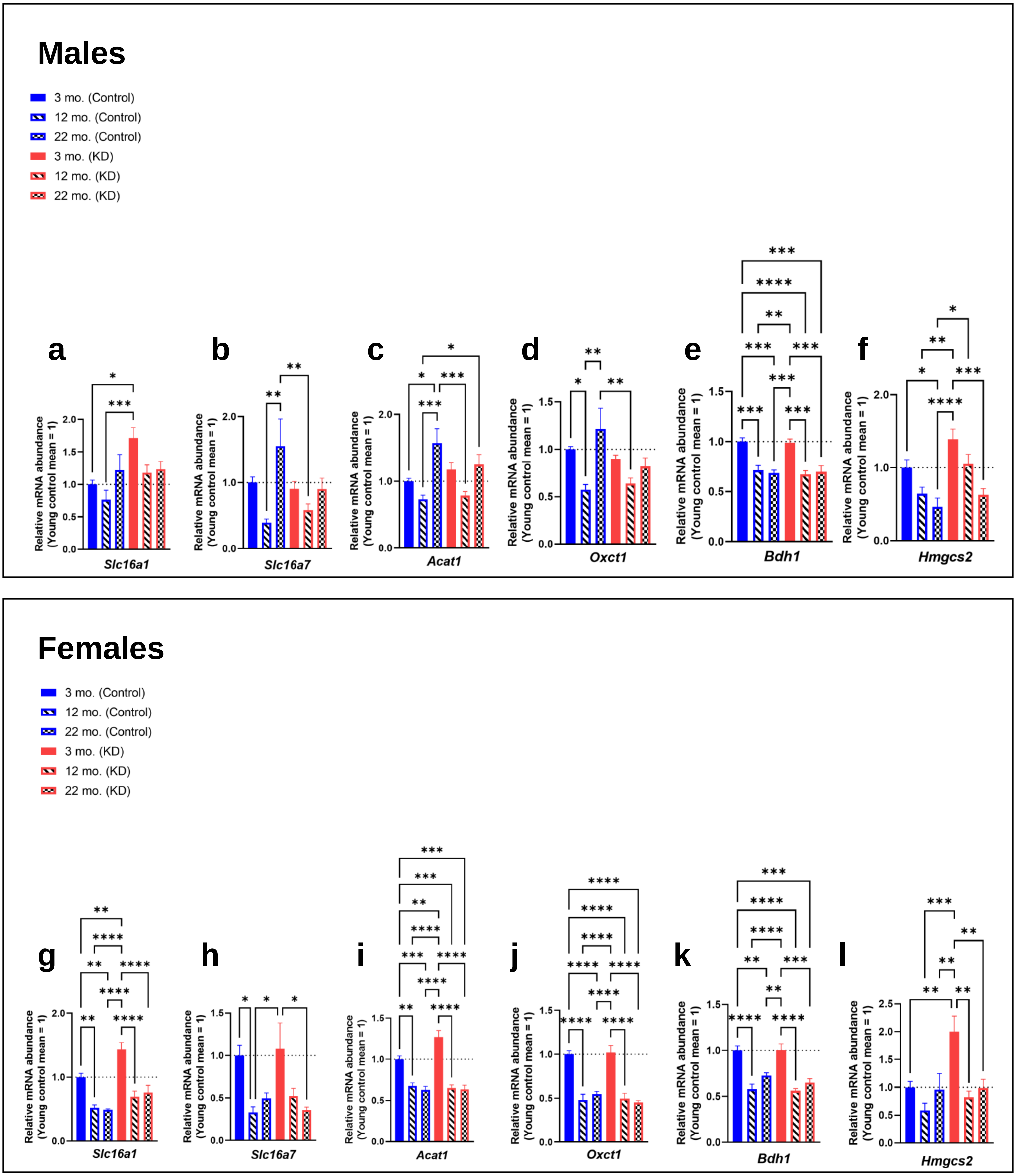


**Extended Data Fig. 8. Gene expression of ketone body metabolism genes in brain.**
**a-f,** *Slc16a1, Slc16a7, Acat1, Oxct1, Bdh1, Hmgcs2* gene expression of day 7 control and KD-fed male mice; n=6-7 per age and diet groups **g-l,** *Slc16a1, Slc16a7, Acat1, Oxct1, Bdh1, Hmgcs2* gene expression of day 7 control and KD-fed female mice; n=6-7 per age and diet groups **a-l,** All data are presented as mean ± SEM; Ordinary one-way ANOVA, Tukey's multiple comparisons test between age and diet groups; *p<0.0332, **p<0.0021, ***p<0.0002. ****p<0.001.

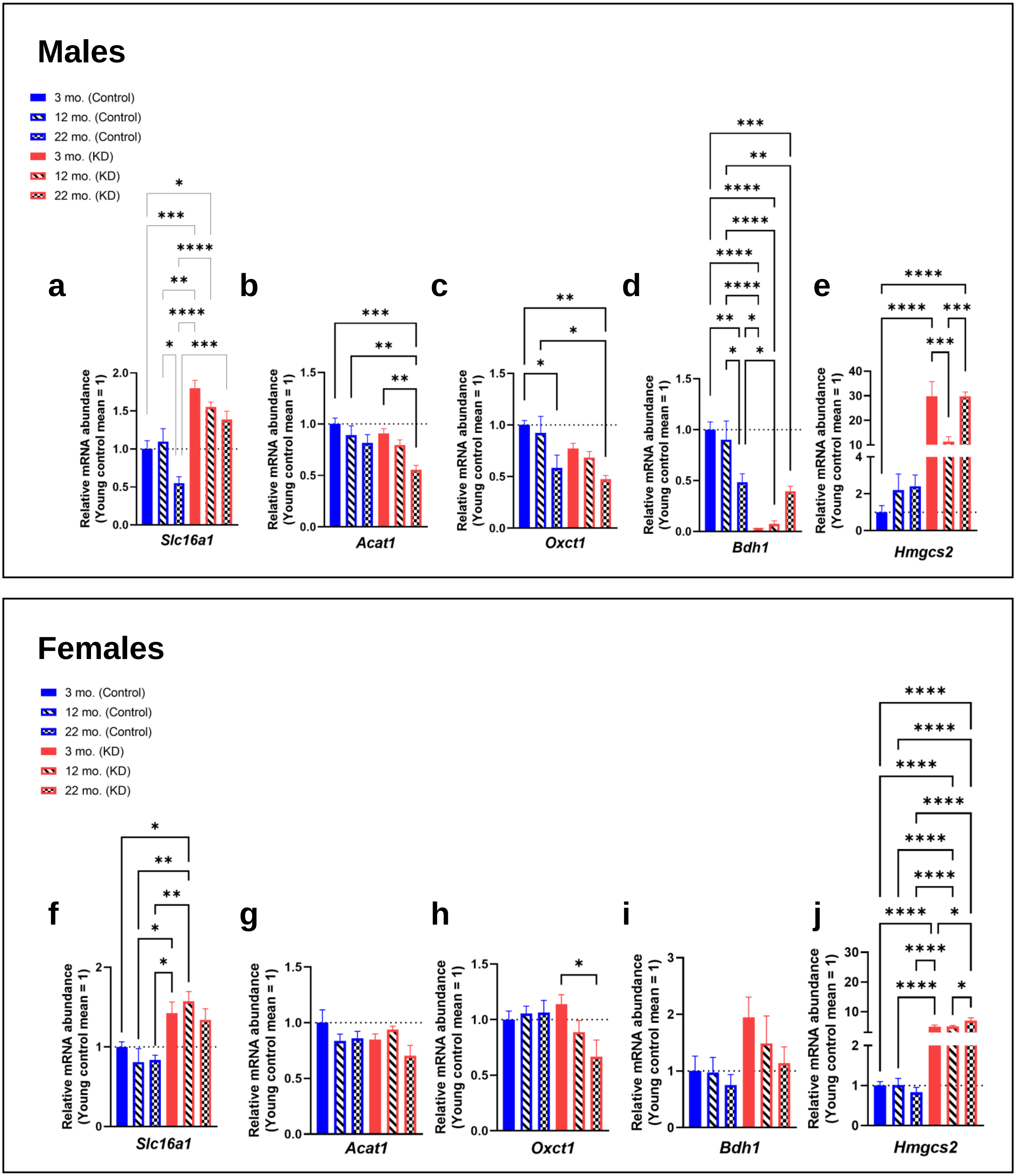


**Extended Data Fig. 9. Gene expression of ketone body metabolism genes in kidney.**
**a-e,** *Slc16a1, Acat1, Oxct1, Bdh1, Hmgcs2* gene expression of day 7 control and KD-fed male mice; n=6-7 per age and diet groups **f-j,** *Slc16a1, Acat1, Oxct1, Bdh1, Hmgcs2* gene expression of day 7 control and KD-fed female mice; n=6-7 per age and diet groups; **a-j;** All data are presented as mean ± SEM; Ordinary one-way ANOVA, Tukey's multiple comparisons test between age and diet groups; *p<0.0332, **p<0.0021, ***p<0.0002. ****p<0.001.

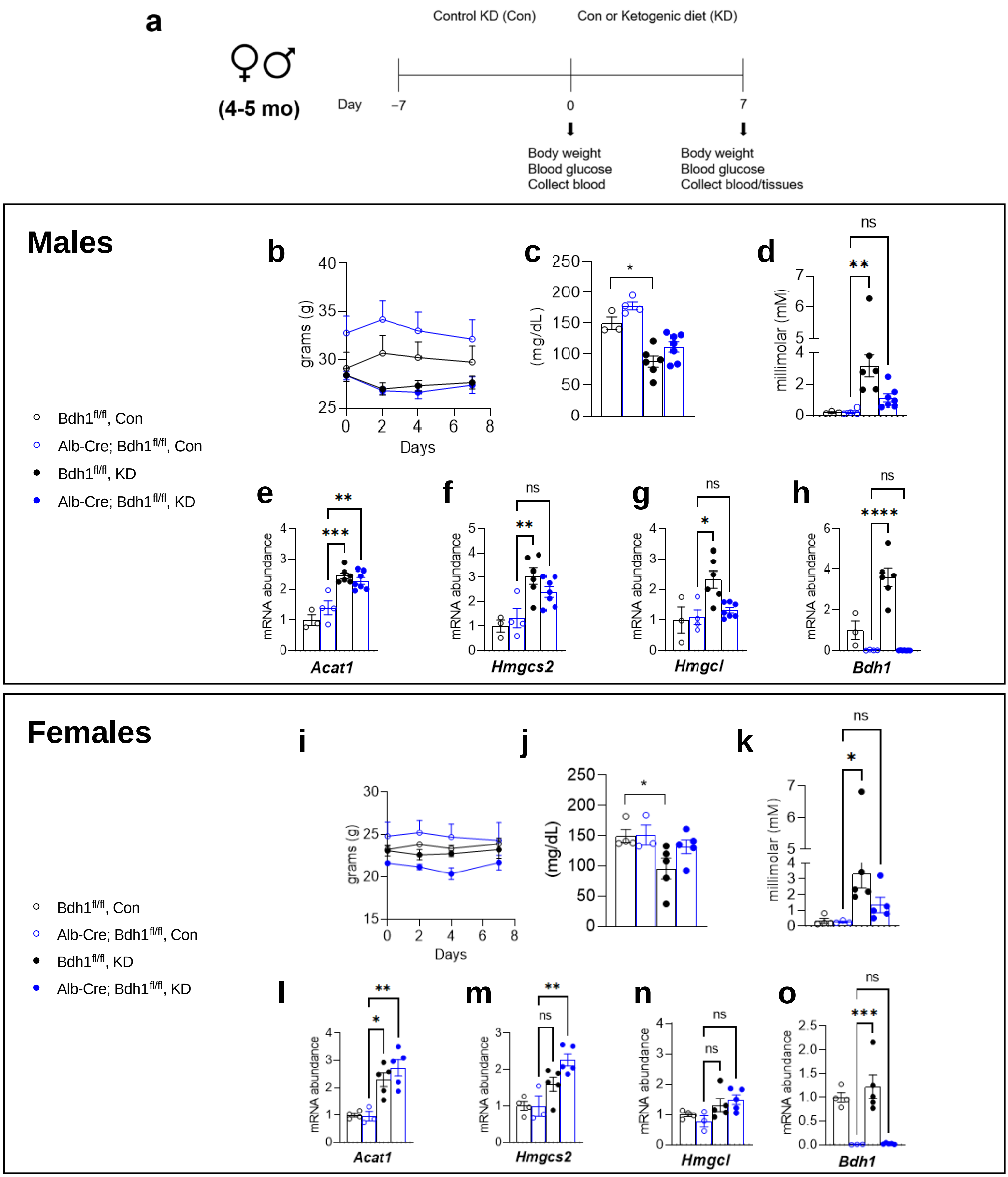


**Extended Data Fig. 10. Male and female *Bdh1* liver knock-out.**
**a,** Experiment and feeding timeline **b,i,** Day 7 body weight (male (top), female (bottom) **c,j,** Day **7** blood glucose (male (top), female (bottom) **d,k,** Day 7 plasma BHB (male (top), female (bottom) **e-h, l-o,** Day 7 *Acat1, Hmgcs2, Hmgcl, Bdh1* gene expression (male (top), female (bottom) **b-o,** *Bdh1* fl/fl control diet, Alb-Cre *Bdh1* fl/fl control diet, *Bdh1* fl/fl KD, and Alb-Cre *Bdh1* fl/fl KD; n=3-7 per group; All data are presented as mean ± SEM; Ordinary one-way ANOVA, Tukey's multiple comparisons test between age and diet groups; *p<0.0332, **p<0.0021, ***p<0.0002. ****p<0.001.
